# Genotypic and phenotypic characterization of a nosocomial outbreak of *Candida auris* in Spain during five years

**DOI:** 10.1101/2024.03.01.582904

**Authors:** Juan Vicente Mulet-Bayona, Irving Cancino-Muñoz, Carme Salvador-García, Nuria Tormo-Palop, Remedios Guna, Fernando González-Candelas, Concepción Gimeno-Cardona

## Abstract

**Objectives:** The investigation of *Candida auris* outbreaks is needed to provide insights into its population structure and transmission dynamics. We genotypically and phenotypically characterized a *C. auris* nosocomial outbreak occurred at the Consorcio Hospital General Universitario de Valencia (CHGUV), Spain.

**Methods:** Data and isolates were collected at CHGUV from September 2017 (first case) until September 2021. Thirty-five isolates, one from an environmental source, were selected for whole genome sequencing (WGS), and the genomes were analyzed along with 335 publicly available genomes, assigning them to one of the five major clades. In order to identify polymorphisms associated with drug resistance, we used the fully susceptible GCA_003014415.1 strain as reference sequence. Known mutations in genes *erg11* and *fks1* conferring resistance to fluconazole and echinocandins, respectively, were investigated. Isolates were classified into aggregating or non-aggregating.

**Results:** All isolates belonged to clade III and were from an outbreak with a single origin. They clustered close to 3 publicly available genomes from a hospital from where the first patient was transferred, being the probable origin. The mutation VF125AL in the *ERG11* protein, conferring resistance to fluconazole, was present in all the isolates and one isolate also carried the mutation S639Y in the *FKS1* protein. All the isolates had a non-aggregating phenotype.

**Conclusions:** Isolates are genotypically related and phenotypically identical but one with resistance to echinocandins, which seems to indicate that they all belong to an outbreak originated from a single isolate, remaining largely invariable over the years. This result stresses the importance of implementing infection control practices as soon as the first case is detected or when a patient is transferred from a setting with known cases.

## Introduction

*Candida auris* was first described in 2009 (1) and rapidly emerged as a serious threat for public health due to its propensity to cause large hospital outbreaks, its tendency to produce bloodstream infections (candidemia) in critically ill patients, its multidrug resistance properties, and its long-term survival in the hospital environment (2). *C. auris* cases have dramatically increased in Europe since 2016 and Spain is one of the countries with the highest prevalence, being responsible of 76% of all the cases and the only European country where candidiasis by *C. auris* is currently considered an endemic situation (3).

Genomic studies, based in whole genome sequencing (WGS) experiments, are increasingly being applied to clinical isolates and are providing insights into how *C. auris* is spreading among facilities, communities, countries and continents (4). To date, five major genetic clades have been described in *C. auris*, separated by many (tens of thousands) single nucleotide polymorphisms (SNPs). Each clade has been associated to a geographical region: clade I (South Asian), clade II (East Asian), clade III (African), clade IV (South American), and clade V (based on a single isolate from Iran separated from the other clades by more than 200,000 SNPs) (4,5).

*C. auris* displays phenotypic characteristics that differ from those of the most common *Candida* species, such as its multidrug resistance profile. Another intriguing characteristic is that two different phenotypes have been described, the aggregating and the non-aggregating, the latter having been linked to a greater virulence of the strains (6).

In September 2017, a *C. auris* outbreak started at the Consorcio Hospital General Universitario de Valencia (CHGUV), Spain, which was reported in previous studies (7–9). Here, we present the results of the WGS analysis and the phenotypic characterization (aggregation and susceptibility) of 35 *C. auris* strains isolated over four years in our setting.

## Methods

### Sample collection

This study is an update to the previous report conducted during 2017-2019 on the same healthcare setting (9). *Candida auris* isolates were recovered from patients admitted to the Consorcio Hospital General Universitario de Valencia (CHGUV), from September 2017 until September 2021. Among all positive cases, 34 *C. auris* isolates sampled during the study period were selected for WGS according to the following criteria: the first two *C. auris* isolates and an echinocandin-resistant isolate from blood were selected, and the remaining isolates were randomly selected from several days apart, attempting to include at least one sample for each sample type and ward where the pathogen had been identified. Furthermore, an isolated environmental sample from a bedside table at the Intensive Care Unit (ICU) was included. All the isolates, previously stored at −70ºC, were subcultured onto Sabouraud Dextrose Agar plates (Becton Dickinson, USA), incubated at 37ºC for 24 h and identified by Matrix-Assisted Laser Desorption/Ionization-Time of Flight (MALDI-TOF, Bruker, USA). The information on the selected isolates is shown in Table 1.

**Table 1.**
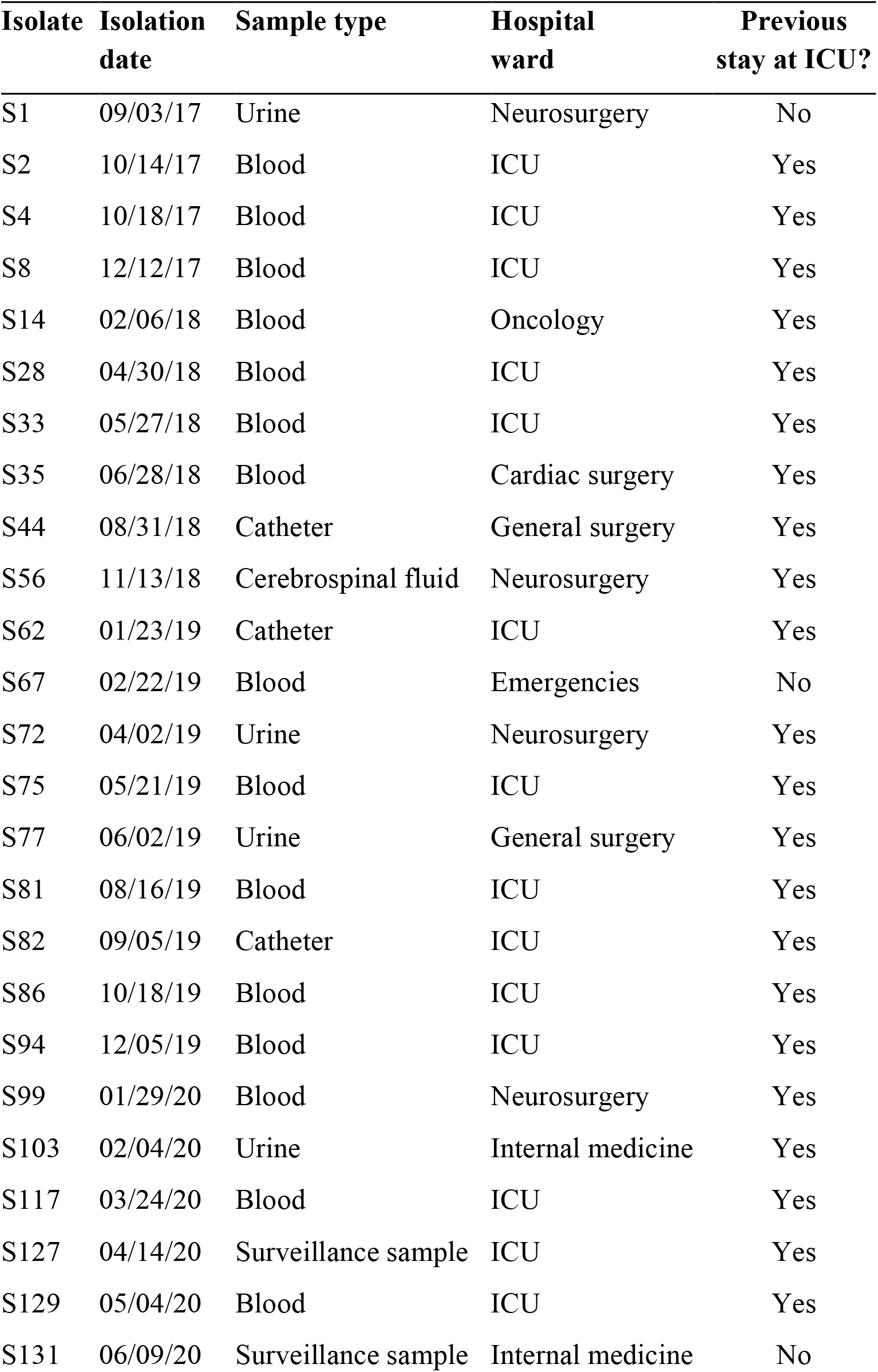
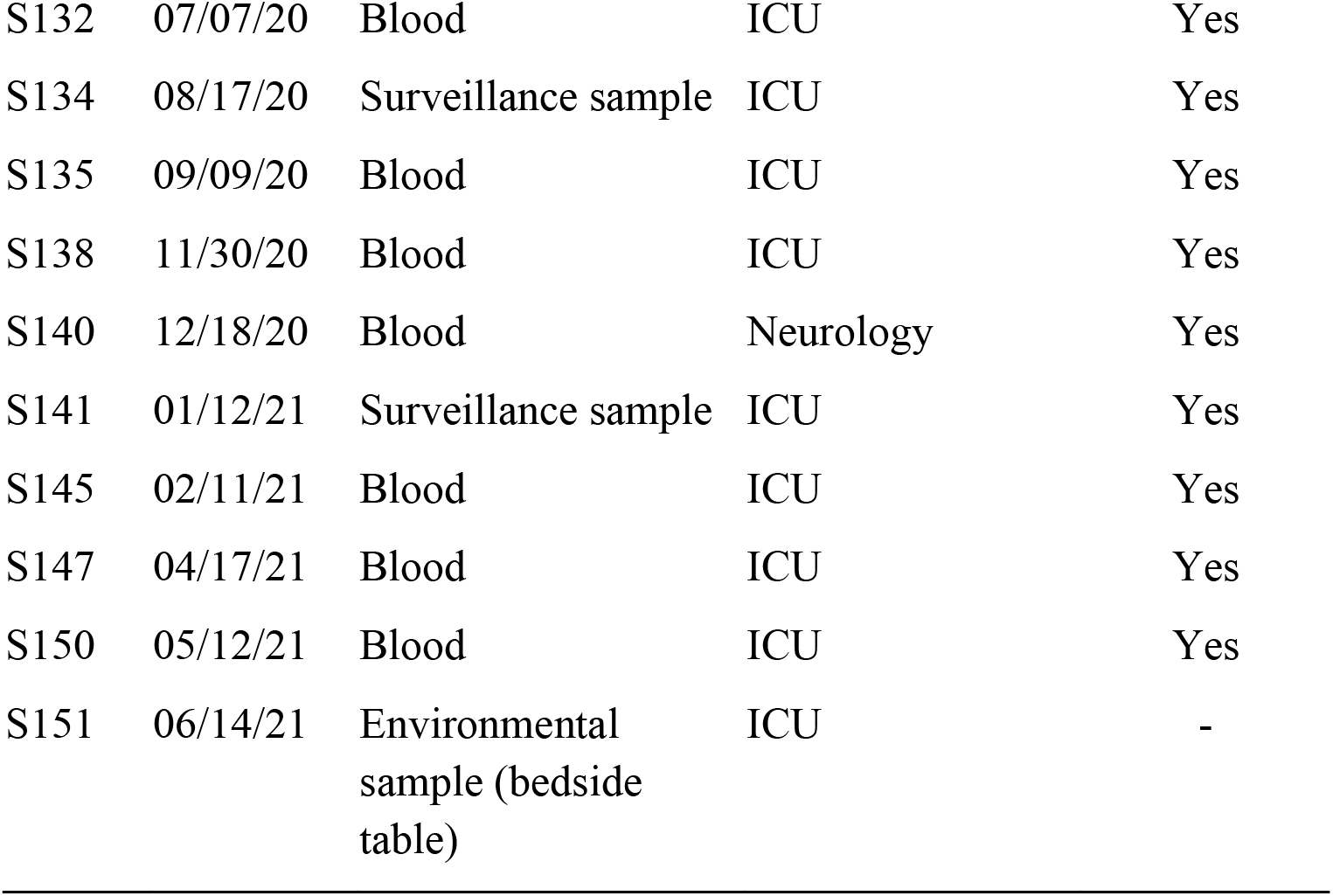
Isolates selected for genotypic and phenotypic characterization.

| **Isolate** | **Isolation date** | **Sample type** | **Hospital ward** | **Previous stay at ICU?** |
| --- | --- | --- | --- | --- |
| S1 | 09/03/17 | Urine | Neurosurgery | No |
| S2 | 10/14/17 | Blood | ICU | Yes |
| S4 | 10/18/17 | Blood | ICU | Yes |
| S8 | 12/12/17 | Blood | ICU | Yes |
| S14 | 02/06/18 | Blood | Oncology | Yes |
| S28 | 04/30/18 | Blood | ICU | Yes |
| S33 | 05/27/18 | Blood | ICU | Yes |
| S35 | 06/28/18 | Blood | Cardiac surgery | Yes |
| S44 | 08/31/18 | Catheter | General surgery | Yes |
| S56 | 11/13/18 | Cerebrospinal fluid | Neurosurgery | Yes |
| S62 | 01/23/19 | Catheter | ICU | Yes |
| S67 | 02/22/19 | Blood | Emergencies | No |
| S72 | 04/02/19 | Urine | Neurosurgery | Yes |
| S75 | 05/21/19 | Blood | ICU | Yes |
| S77 | 06/02/19 | Urine | General surgery | Yes |
| S81 | 08/16/19 | Blood | ICU | Yes |
| S82 | 09/05/19 | Catheter | ICU | Yes |
| S86 | 10/18/19 | Blood | ICU | Yes |
| S94 | 12/05/19 | Blood | ICU | Yes |
| S99 | 01/29/20 | Blood | Neurosurgery | Yes |
| S103 | 02/04/20 | Urine | Internal medicine | Yes |
| S117 | 03/24/20 | Blood | ICU | Yes |
| S127 | 04/14/20 | Surveillance sample | ICU | Yes |
| S129 | 05/04/20 | Blood | ICU | Yes |
| S131 | 06/09/20 | Surveillance sample | Internal medicine | No |
| S132 | 07/07/20 | Blood | ICU | Yes |
| S134 | 08/17/20 | Surveillance sample | ICU | Yes |
| S135 | 09/09/20 | Blood | ICU | Yes |
| S138 | 11/30/20 | Blood | ICU | Yes |
| S140 | 12/18/20 | Blood | Neurology | Yes |
| S141 | 01/12/21 | Surveillance sample | ICU | Yes |
| S145 | 02/11/21 | Blood | ICU | Yes |
| S147 | 04/17/21 | Blood | ICU | Yes |
| S150 | 05/12/21 | Blood | ICU | Yes |
| S151 | 06/14/21 | Environmental sample (bedside table) | ICU | - |

### Antifungal susceptibility testing (AFST) and phenotypic characterization

AFST was carried out in 716 isolates, in the context of the daily laboratory routine, by broth microdilution with either VITEK^®^ 2 AST-YS08 (bioMérieux, France) or Sensititre^®^ YeastOne YO10/ITAMYUCC (Thermo Fisher Scientic, USA). Tentative breakpoints from CDC were used to describe the susceptibility of the isolates (10). The selected isolates (n=35) were classified as aggregating or non-aggregating, as described by Szekely *et al*. (11). In all cases, Type strain MYA-5001^®^ was used as a control.

### DNA extraction and whole genome sequencing

Several colonies of each isolate were introduced in Eppendorf tubes with distilled water and centrifuged at 5,000 rpm for 5 minutes. The supernatant was removed, 180 μL of lysis buffer (ATL buffer; Qiagen) and 40 μL of proteinase K (Qiagen) were added, and the mix was incubated at 56 ºC overnight. Then, DNA was extracted from the mix using the QIAsymphony^®^ SP DNA kit for the QIAsymphony^®^ SP system (Qiagen). Extracted DNA was quantified using Tecan Infinite^®^ PRO fluorometer (Tecan, Switzerland) and the concentrations were adjusted to 5 ng/μL.

DNA libraries were prepared with the Illumina DNA Prep kit (Illumina) following the Illumina DNA Prep Reference Guide v10. Input DNA was diluted to 0.2 ng/μL to start the protocol. At the multiplexing step, Nextera DNA CD Indexes (Illumina) were used. Library size selection was performed according to the Illumina library Prep protocol with Sample Purification Beads (SPB) included in the library Prep kit. Library size was validated by using Fragment Analyzer 48-Capillary Array with the HS NGS Fragment Kit (1-6000bp) (Agilent). Libraries were sequenced with a Nextseq 2×150 pb paired-end reagent kit (NextSeq 500/550 High Output Kit v2.5, Illumina) on an Illumina Nextseq500 sequencer, according to manufacturer’s instructions.

### Bioinformatic analysis

In order to place our samples in a global phylogenetic context, we downloaded 334 publicly available *C. auris* genomes from all the continents, including the reference assemblies for all the major clades (GCA_002759435.2 for Clade I, GCA_003013715.2 for Clade II, GCF_002775015.1 for Clade III, and GCA_003014415.1 for Clade IV, and strain IFRC2087 for clade V (12)). Based on these assemblies, sequencing reads were simulated and created using the ART program (13). All the information regarding the 369 analyzed genomes is available in Supplementary Table 1.

Raw sequencing reads were trimmed and filtered using fastp (14) with the following parameters “--cut_tail, --cut_window_size=10, --cut_mean_quality=20, --length_required=50, --correction, and – dedup”. Variant calling was carried out using snippy software (https://github.com/tseemann/snippy) with the parameters “--mincov 10 --minfrac 0.9 --minqual 60”. Raw sequences were deposited in the European Nucleotide Archive server under Bioproject PRJEB70513.

### Sequence alignment and phylogenetic analysis

Different SNP-based multiple sequence alignments (MSAs) were constructed depending on the analytical purpose. First, an MSA including all previously described global genomes (n=369) was constructed to put our samples into a global phylogenetic context. The second MSA was built using only genomes belonging to Clade III (n=268), including samples from two previous nosocomial outbreaks (15,16). Lastly, a MSA which included those isolates that were genetically close to our outbreak (n=40) was constructed. These MSAs consisted of 190,611, 803, and 105 variable positions, respectively.

MSAs were constructed from the SNPs detected by mapping against the reference genome of *C. auris* clade III (GCF_002775015.1). Those polymorphisms were called using the Snippy program (https://github.com/tseemann/snippy) with a minimum coverage of 10x and a minimum frequency of 90%. The detected SNPs were then concatenated using the “snippy-core” option in Snippy and masked to those polymorphisms detected in duplicated and repeated regions of the reference genome (approx. 3.2% of the entire genome). Pairwise SNP distances were calculated with the R package ape.

All maximum-likelihood phylogenetic trees were constructed using IQ-TREE2 (17) using the GTR substitution model with 1,000 bootstrap replicates, and considering the invariant sites of the whole genome alignment.

### Identification of antifungal resistance mutations

In order to identify polymorphisms associated with drug resistance, we used the fully susceptible GCA_003014415.1 strain as reference sequence. SNPs were annotated using snpEff (18). Known mutations in genes *erg11* and *fks1* were detected for fluconazole and echinocandins, respectively.

## Results

### Description of the outbreak

The first *C. auris* in our setting was isolated in September 2017, in a urine sample of a patient transferred from a hospital located in the same city (Hospital Universitari i Politècnic La Fe; HUiP La Fe, Valencia, Spain). HUiP La Fe had recently reported four *C. auris* candidemia cases, which later also became an outbreak (19,20). One month later, *C. auris* was isolated in a blood culture from another patient hospitalized in the Intensive Care Unit (ICU), therefore being the first candidemia case. Since then, candidemia cases have kept being reported, in spite of having implemented the recommended infection control practices, such as isolation/cohorting of patients, enhanced cleaning with chlorine-based disinfectants or screening studies for detecting colonized patients (21). The screening studies were implemented to ICU patients, at admission and once a week, and consisted of culturing axillary-rectal and pharyngeal swabs. Until September 2021 (four years of follow-up), the outbreak affected 550 patients, considering both colonized and infected patients. The most frequent infection was candidemia, with 86 cases. The temporal distribution of *C. auris* colonization or infection episodes is shown in Figure 1.

**Figure 1.**
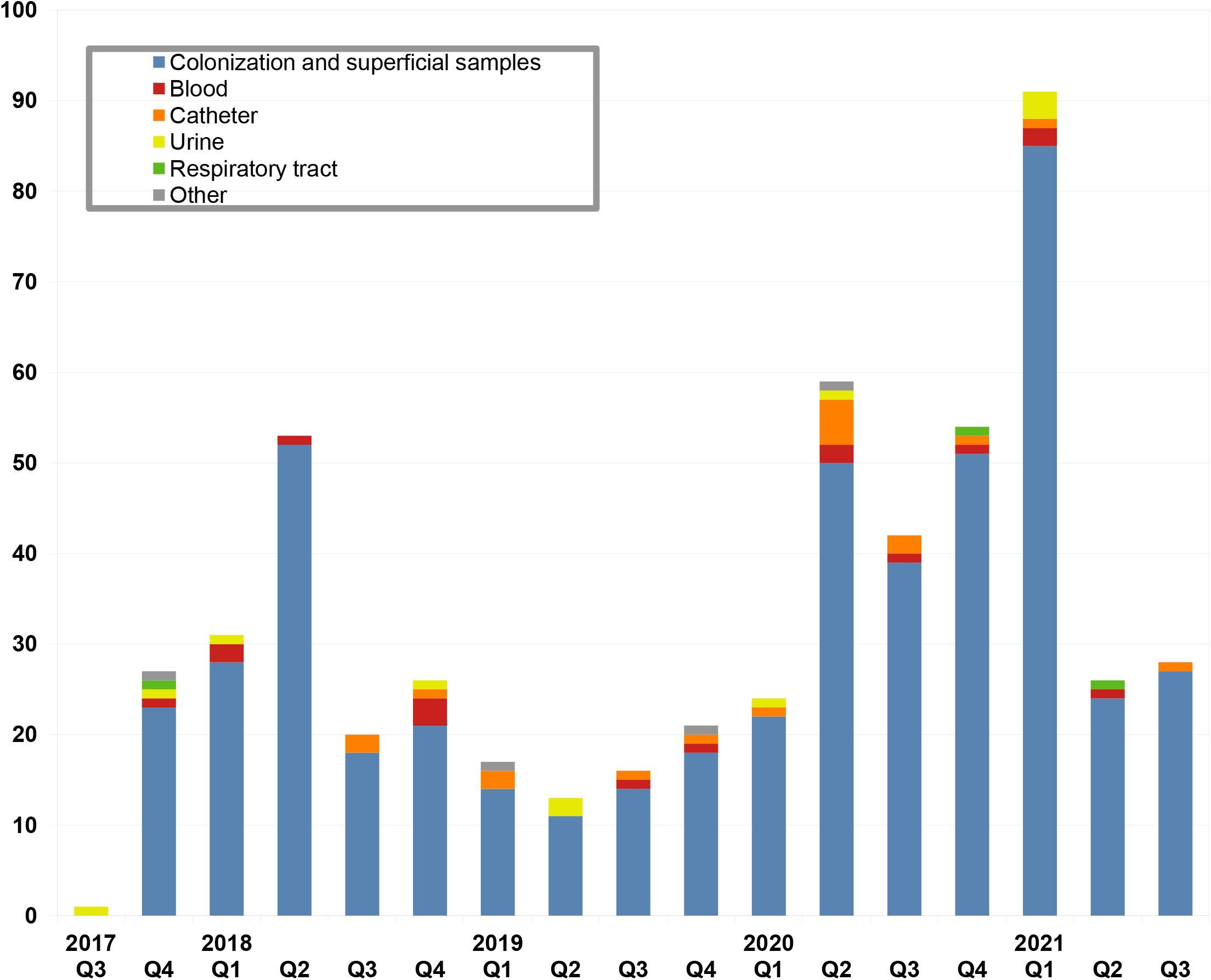
First-occurring episodes of *C. auris* colonization or infection (n=550 patients) from September 2017 (first case) until September 2021. Q: calendar quarter. Isolates from every calendar quarter have been selected for sequencing.

The pathogen has been identified among individuals admitted in nearly all wards within the facility, although the ICU was by far the most frequent unit, with 468 affected patients (including surveillance samples), followed by the Internal Medicine/Infectious Diseases Unit with 12 patients. Therefore, the outbreak seems to be established at the ICU, being the main epidemiological source. It is noteworthy that the outbreak worsened from June 2020 to February 2021, due to the COVID19 pandemic, probably due to the over-occupancy of the ICU together with the higher workload of healthcare workers (8). The thirty-day crude mortality rate for candidemia cases was 32.6% (9).

### Phenotypic and genotypic characterization of the outbreak

Regarding AFST, all the isolates (n=716) were resistant to fluconazole, 2.8% were also resistant to echinocandins and 0.6% to amphotericin B. However, most echinocandin or amphotericin B isolates were from colonization samples and only one echinocandin resistant isolate was from an invasive sample (blood). This isolate developed resistance to echinocandins during a long treatment with echinocandins (22). All the isolates were phenotypically characterized as non-aggregating (Figure 2).

**Figure 2.**
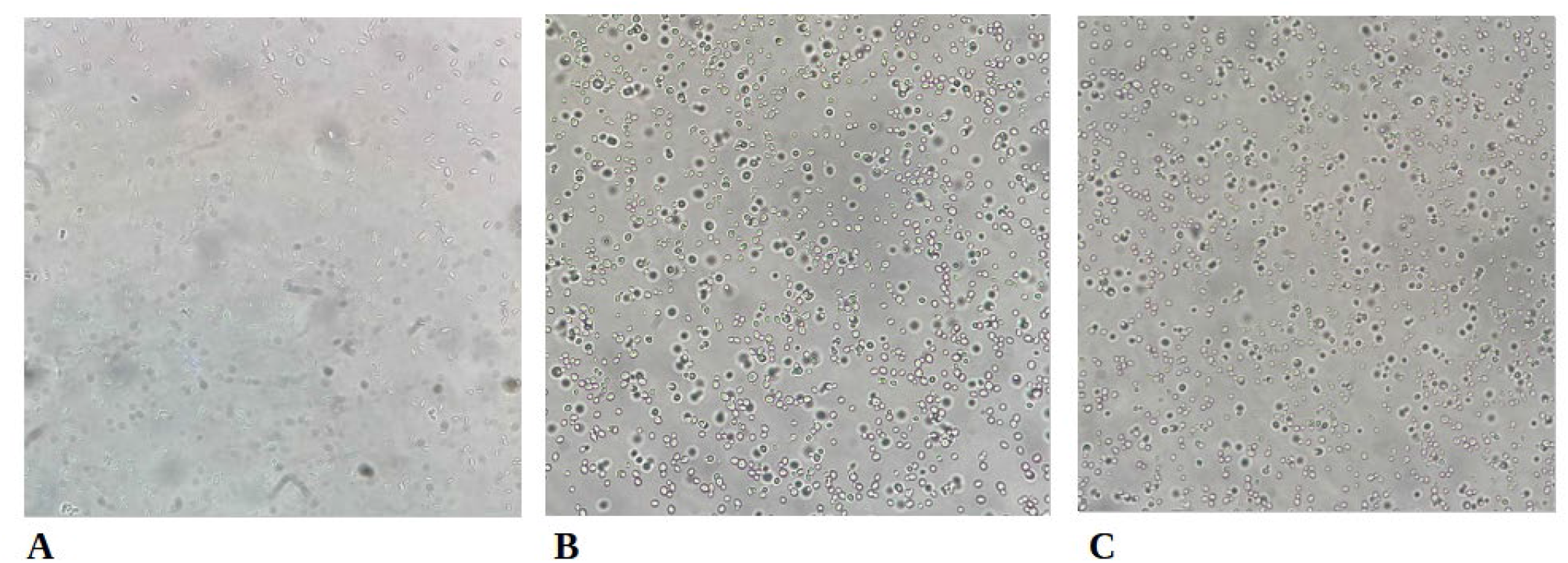
Microscopic appearance of isolates MYA-5001 (type strain, **A**), S1 (**B**) and S151 (**C**), they all showing disperse cells and, therefore, being characterized as non-aggregating (x400 magnification). All thirty-five isolates showed the same non-aggregating phenotype.

All the sequenced isolates carried the VF125AL mutation (also referred to as F126L) in the *ERG11* protein associated with fluconazole resistance. Moreover, one also harbored the S639Y mutation related to echinocandin resistance in the *FKS1* protein, being consistent with the AFST profile (Supplementary Table 1).

### Phylogenetic analysis of the outbreak

The global phylogenetic analysis of the 35 *C. auris* genomes obtained (Supplementary Figure 1) showed that all belong to Clade III (South African). Given that the median genetic distance between the samples of our outbreak and the other clades was very large (Clade I, 21,469 SNPs; Clade II, 30,916 SNPs; Clade IV, 79,368 SNPs; and Clade V, 126,769 SNPs), we decided to zoom in on the samples genetically most closely related to it. The phylogenetic analysis of this dataset revealed four isolates genetically and temporally related to our outbreak isolates, with a median distance of 5 SNPs (range 1-9 SNPs). Three of these were from 2016 and another hospital (HUiP La Fe) located in the same city as our setting (strains “AA-194”, “AA-200” and “AA-214”), and were phylogenetically ancestors of our isolates. Interestingly, a case from Austria reported in 2021 (23) was also genetically related to our outbreak with a median distance of 6 SNPs (Strain “Cau4”) (Figure 3). Based on the epidemiological information of this case, the patient had been admitted to a hospital in Valencia, so it is very likely that the infection occurred during that period.

**Figure 3.**
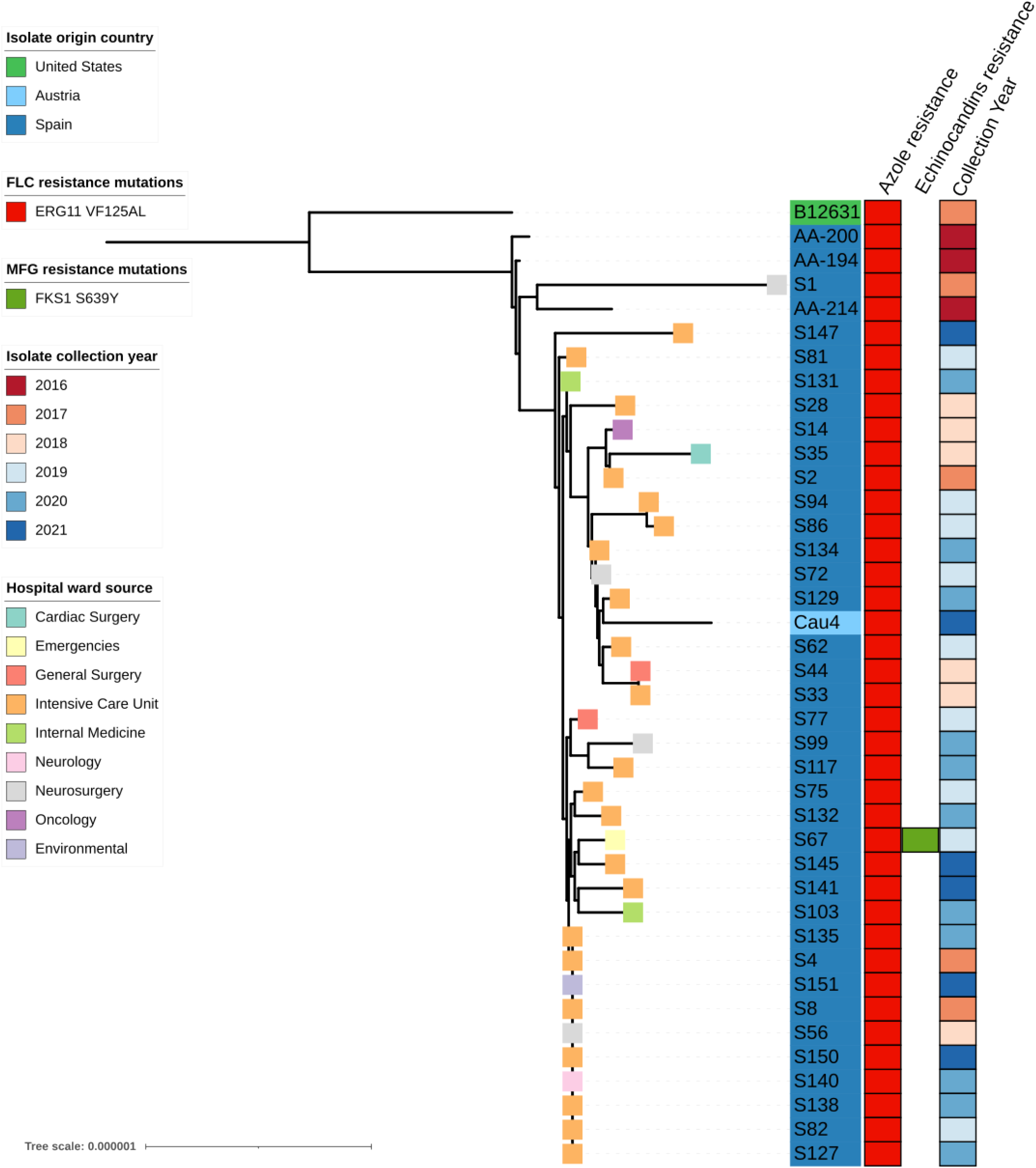
Phylogeny of the Valencia outbreak. The tree was contructed using the 35 *C. auris* genomes from our setting (strains from “S1” to “S151”) and five publicly available genomes. The strain “B12631” was used as an outgroup. Country of origin, azole- and echinocandin-resistance associated mutations, and collection year are shown. The rectangle shape at the branch tip indicates the hospital ward origin.

The population structure of the outbreak suggests that the pathogen had a single common source (monophyletic cluster), probably originating in 2016 in another regional hospital (HUiP La Fe). From there, the pathogen was introduced to our hospital in September 2017 by a patient referred from HUiP La Fe (“S1” strain) and from there it has dispersed throughout the hospital during the study period. The maximum genetic distance within the outbreak is 12 SNPs, suggesting recent transmissions within an ongoing outbreak (24) (Figure 4). Interestingly, we found that the most closely related genome to our outbreak was an isolate from Indiana, USA (strain “B12631”). However, because of the median genetic distance observed (26 SNPs, range 22-36), and because we found no epidemiological link between this strain and the rest of the outbreak, the idea of direct transmission was discarded.

**Figure 4.**
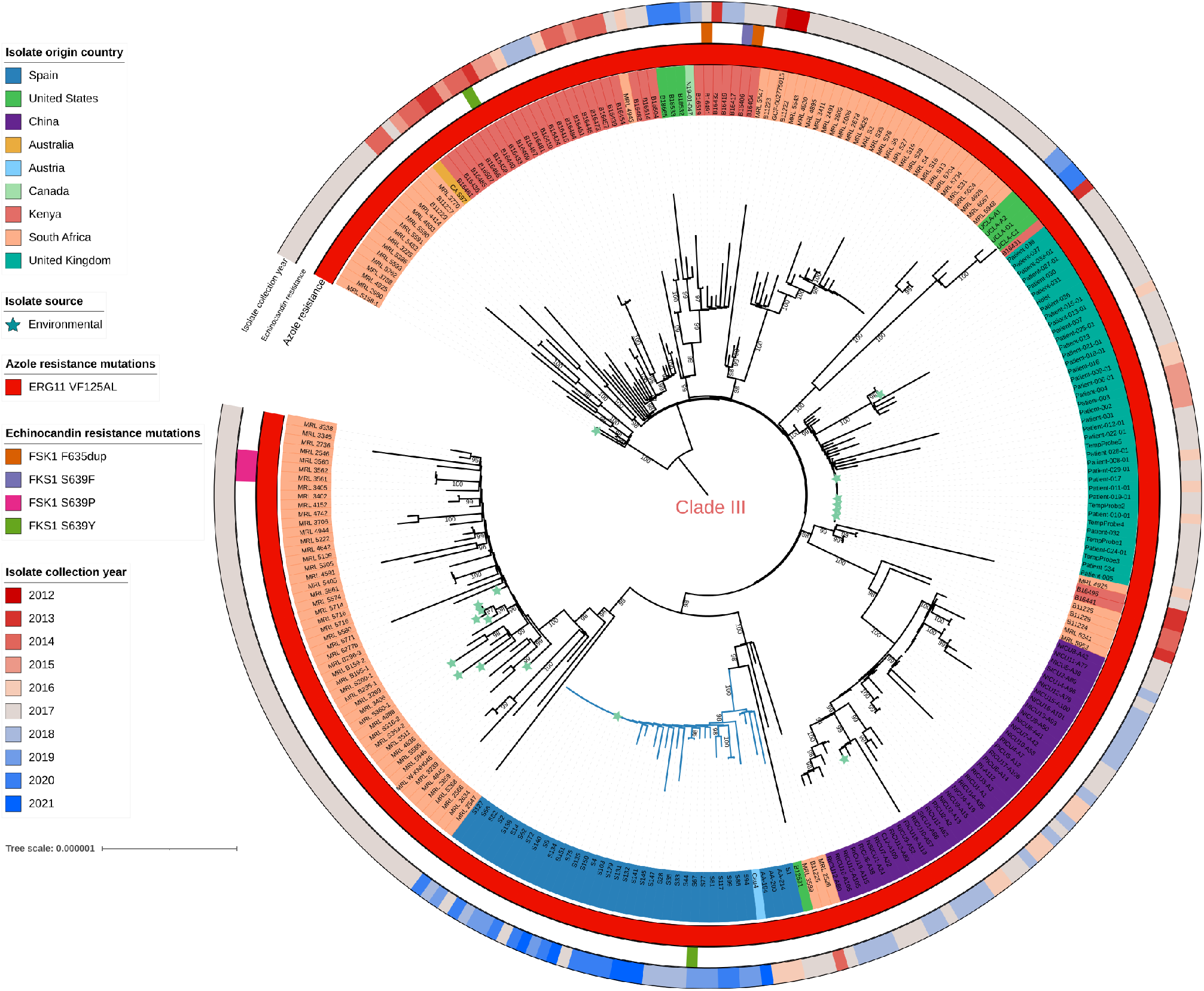
Maximum likelihood phylogenetic tree of 35 *C. auris* genomes from our setting (S1 to S151) and 268 publicly available *C. auris* genomes from Clade III (n= 803 variant positions). Country of origin, azole- and echinocandin-resistance associated mutations, and collection year are shown from inner to outer circles. The star shape indicates isolates of environmental origin.

**Figure 5.**
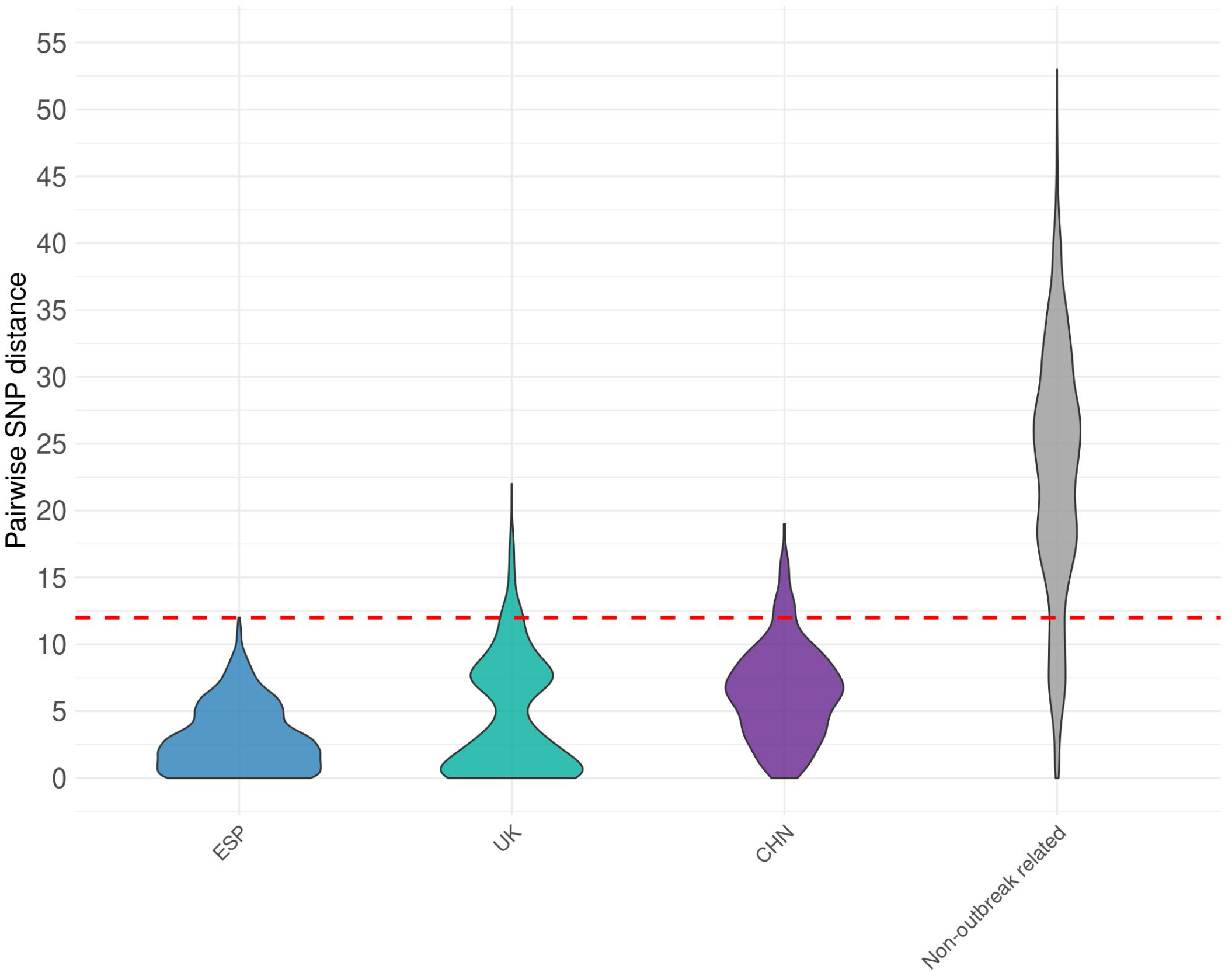
Pairwise genetic distances between samples belonging to the same nosocomial outbreak (intra-outbreak) and those that are not considered to be from the same outbreak. The red dashed line indicates the threshold marker of recent transmission (12 SNPs).

### Determining transmission

By analyzing the genetic distance as a marker of recent transmission within the same facility (≤12 SNPs), we observed that our outbreak fell within this threshold; however, this approximation did not fit with other nosocomial outbreaks. Examining the median genetic distance within other previously described outbreaks (one from the UK (15) and one from China (16)), we observed that the maximum distance was 22 SNPs for the case of the UK outbreak, and 19 SNPs for the China outbreak (Figure 4). In contrast, when analyzing the range of distances between the rest of the genomes that have not been described in nosocomial outbreaks, we can observe that there are pairs of isolates that have compatible distances with this category of recent transmission (range 0-53 SNPs). This, along with the fact that the first reported case was a patient transferred from another hospital, suggests that the threshold for recent transmission may have to be redefined.

## Discussion

*C. auris* outbreaks have been difficult to control worldwide, with some of them lasting for over a year (20,25,26), as the one occurred in our setting. The characterization of the isolates show that they are genotypically related and phenotypically non-aggregating, which seems to indicate that they all belong to a single outbreak originated from a single isolate, remaining largely invariable over the years. It is noteworthy that the first patient was not admitted to the ICU, while the outbreak started one month later in the ICU ward; no epidemiological relationship was found. Since then, the pathogen has affected mostly patients at the ICU or recently discharged from it, being the probable main epidemiological source. However, other wards cannot be excluded as epidemiological sources, as screening colonization studies were performed only at the ICU and there may be undetected colonized patients in other wards, who do not develop as many invasive infections due to less risk factors. Environmental screening was also performed in some cases at the ICU, finding the pathogen in beds, bed rails, bedside tables, computer keyboards and mouses, and several medical devices. In fact, it is widely recognized that invasive candidiasis, such a *C. auris* candidemia, has a propensity to affect critically ill patients in ICUs (20,25,27).

Some *C. auris* outbreaks have been attributed to recent spread to other continents, as the first European outbreak occurred in United Kingdom in 2015 (25), whose isolates analyzed by WGS were found to belong to clade I (with an Asian origin). *C. auris* was isolated for the first time in our setting in September 2017, in a patient transferred from HUiP La Fe, which had an ongoing outbreak (19,20). Amplified fragment length polymorphism (AFLP) analysis in samples from HUiP La Fe revealed that the isolates were genetically similar to South African isolates (clade III), except one isolate which was grouped in the Venezuelan cluster (clade IV) (20). Three isolates from this outbreak were also sequenced (isolates AA-194, AA-200 and AA-214) and assigned to clade III (4). Although the most likely hypothesis was that our outbreak originated from the outbreak of HUiP La Fe, the possibility that *C. auris* would have been introduced to our setting from other source did exist. In fact, Chow et al. found that multiple clades of *C. auris* were introduced into the United States, some even several times (24). However, the phylogenetic analysis of the 35 isolates from the CHGUV showed that they all belonged to clade III, and clustered with the 3 isolates from HUiP La Fe, which were also ancestors of our isolates, making the first hypothesis more likely. Spanish isolates had a maximum distance of 12 SNPs among them, suggesting that these two outbreaks have evolved locally from a single origin (Figure 3). Previous studies have used a maximum of 12 SNPs among isolates to delimit an outbreak in a single hospital (24,28). However, this is not necessarily a universal limit, applicable in all cases. By analyzing the genetic distance among samples from the same nosocomial outbreak in two different clinical sites, we have observed that it is higher (Figure 4). This variation may be due to the pathogen not evolving at the same rate in all cases, or to slight differences in the pattern of transmission among settings. This might be reflected in different topological patterns in the corresponding phylogenetic trees. For instance, the establishment of single strains in different locations in the same facility might lead to sustained divergence among them, similar to allopatric differentiation in incipient speciation. In fact, these topological patterns seem to be present in the cases of the China outbreak, where we can observe the generation of different clades from the same origin (Figure 4). More detailed information about the time and location of the isolates within the hospital along with matching environmental samples would be necessary to test this hypothesis. Interestingly, a case from Austria (strain “Cau4”) was found to be closely related to the Spanish isolates. Using public epidemiological information on the isolate, we realized that the patient had been admitted to a hospital in Valencia during October 2021, suggesting that the infection occurred during his time in the facility (23). The maximum distance between Spanish and other clade III isolates was 60 SNPs, being in line with the mean number of SNPs within each cluster, as reported previously (<70 SNPs) (4).

Genome studies showed that several known antifungal drug targets for *C. albicans* are conserved in *C. auris*, including the azole target lanosterol 14 α-demethylase (*ERG11*) and the echinocandin target 1,3-beta-glucan synthase (*FKS1*), and several mutations in those genes have been described and linked to drug resistance, in both *C. albicans* and *C. auris* (4,29). The mutation VF125AL (also referred to as F126) in *ERG11* was present in all isolates from our setting, conferring them resistance to azole drugs, which was consistent with their phenotypic behavior. This mutation is strongly associated with the geographic clade III, as it has been reported in clonal clade III isolates from South Africa (29). Our isolates belong to group G3 of clade III, only integrated by fluconazole-resistant isolates with the mentioned mutation, while groups G1 and G2 are integrated by isolates without fluconazole-resistance associated mutations. With respect to echinocandins, several hot-spot mutations in *FKS1* have been reported to cause resistance to these antifungals (in positions F635, S639 and R1354) (29–32). Mutation S639Y was found in the echinocandin-resistant isolate S67, which was isolated from a patient carrying a long-term peripherally inserted central catheter line and who was previously treated with echinocandins, probably selecting the raise of resistance mutants (this case was previously reported in (33)). The S639Y mutation has been reported to cause echinocandin resistance in some *C. auris* isolates from clades I and III (31,32).

Two different phenotypes have been described for *C. auris:* the aggregating and the non-aggregating phenotypes, the latter having been linked to a greater virulence of the strains (6). However, the genetic regulation of the *C. auris* morphogenesis and its *in vivo* significance remains largely uncharacterized, although some insights into the *C. auris* genome have been provided (34). The crude mortality rate for candidemia cases in our setting (32.6%) is in line with the mortality reported from other outbreaks, from 20% to 60%, so it is difficult to evaluate if the non-aggregating phenotype of our isolates has contributed to a higher mortality. Instead, Sherry et al. found that the non-aggregating phenotype strains produced more biofilm (35), so this fact might explain the high prevalence of invasive infections (candidemia) and the difficulty in ending the outbreak in our setting, as biofilm-producing bacteria have a better adherence to medical devices and surfaces. In a previous study, isolates belonging to different clades showed different phenotypic traits, including the aggregating/non-aggregating phenotype, being therefore possible that isolates from every clade had an associated phenotypic behavior (11). Interestingly, all our clade III isolates exhibited a non-aggregating phenotype, differing from the clade III isolates from Szekely et al., which were all strongly-aggregating isolates (11). Therefore, it seems that phenotypic traits are not uniform within a clade, but they are strain-dependent and remain constant over the years. More isolates from different clades and countries should be tested in order to better assess the heterogeneity and the clinical significance of the phenotype.

A limitation of this study is that only a small subset of isolates among all positive cultures has been characterized, so a different strain might have been missed. However, the random selection from several days apart over four years should minimize that risk. The molecular epidemiological investigation of *C. auris* outbreaks from different locations is clearly needed to understand the population structure and transmission dynamics of this novel pathogen. Our results provide evidence that nosocomial outbreaks are usually originated from a single origin. Patient environments serve as reservoirs of the pathogen and this results in a cross-transmission among patients within the hospital setting, usually in ICUs. This fact should stress the importance of implementing infection control practices as soon as the first case is detected or when a patient is transferred from a setting with known cases.

## Acknowledgements

Irving Cancino Muñoz was supported by a Margarita Salas contract (UNI/551/2021) from Ministerio de Universidades (Spanish Government). This work was supported by projects CIPROM-2021-053 from Generalitat Valenciana and PID2021-127010OB-I00 from Ministerio de Ciencia e Investigación (Spanish Government). Sequencing was funded by a Dr. López Trigo research promotion award, granted to Juan Vicente Mulet-Bayona by the Fundación de Investigación del Hospital General Universitario de Valencia.

---

**Supplementary Figure 1.**
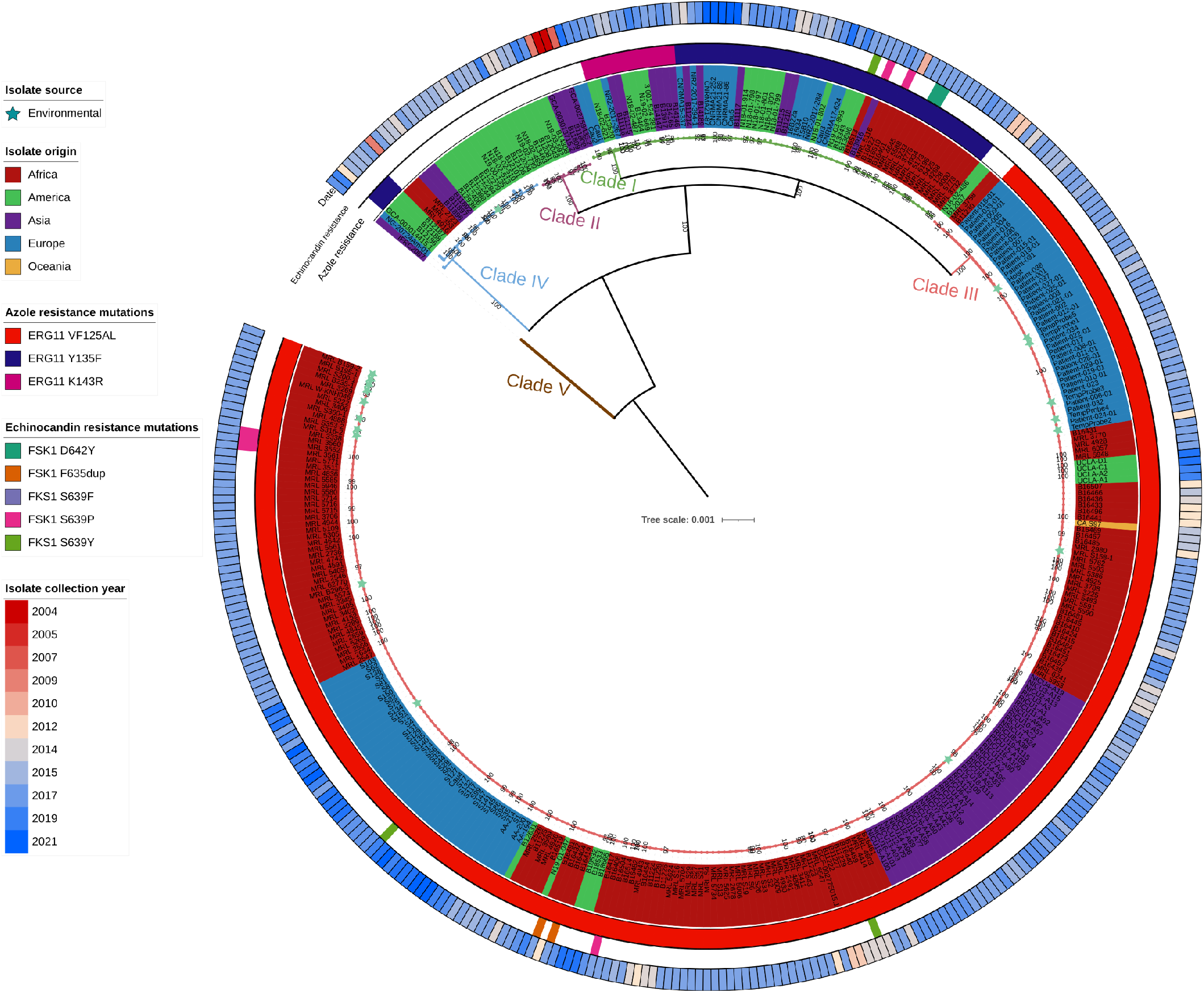
Maximum likelihood phylogenetic tree of 35 *C. auris* genomes from our setting (S1 to S151) and 334 publicly available *C. auris* genomes (n= 190,611 variant positions). Geographic clades, country of origin, azole- and echinocandin-resistance associated mutations, and collection year are shown from inner to outer circles. The star shape indicates isolates of environmental origin.

